# Wnt/β-catenin signaling is required for the development of multiple nephron segments

**DOI:** 10.1101/488718

**Authors:** Patrick Deacon, Charles W. Concodora, Eunah Chung, Joo-Seop Park

## Abstract

The nephron is composed of distinct segments that perform unique physiological functions to generate urine. Little is known about how multipotent nephron progenitor cells differentiate into different nephron segments. It is well known that Wnt/β-catenin signaling regulates the maintenance and commitment of mesenchymal nephron progenitors during kidney development. However, it is not fully understood how it regulates nephron patterning after nephron progenitors undergo mesenchymal-to-epithelial transition. To address this, we performed β-catenin loss-of-function and gain-of-function studies in epithelial nephron progenitors in the mouse kidney. Consistent with a previous report, the formation of the renal corpuscle was defective in the absence of β-catenin. Interestingly, we found that epithelial nephron progenitors lacking β-catenin were able to form presumptive proximal tubules but that they failed to further develop into differentiated proximal tubules, suggesting that Wnt/β-catenin signaling plays a critical role in proximal tubule development. We also found that epithelial nephron progenitors lacking β-catenin failed to form the distal tubules. Constitutive activation of Wnt/β-catenin signaling blocked the proper formation of all nephron segments, suggesting tight regulation of Wnt/β-catenin signaling during nephron patterning. This work shows that Wnt/β-catenin signaling regulates the patterning of multiple nephron segments along the proximo-distal axis of the mammalian nephron.

## Introduction

The nephron is composed of multiple segments (Desgrange and Cereghini, 2015). Podocytes and the Bowman’s capsule in the renal corpuscle are followed by the proximal tubule (PT), loop of Henle (LOH), and distal tubule (DT) (Desgrange and Cereghini, 2015). Each of these segments performs unique physiological functions, indicating that different cell types are present in different nephron segments (Adam et al., 2017; Lee et al., 2015). All nephron tubule cells are derived from the multipotent mesenchymal nephron progenitors (MNPs) which are located at the cortex of the developing kidney (Boyle et al., 2008; Kobayashi et al., 2008). When a tip of the collecting duct (CD) branches, a subset of MNPs located underneath the tip undergoes mesenchymal-to-epithelial transition, forming the renal vesicle (RV). Epithelial nephron progenitors in the RV undergo complex morphogenesis to form the S-shaped body (SSB) which eventually develops into a nephron (McMahon, 2016).

Little is known about which signaling pathways direct MNPs to develop into distinct nephron segments. It had been reported that, in the mouse kidney, Notch signaling promotes the formation of PT and represses the formation of DT (Cheng et al., 2007; Cheng et al., 2003; Surendran et al., 2010). However, we have recently shown that Notch signaling regulates the formation of all nephron segments in the mouse kidney without preferentially promoting the formation of a specific segment (Chung et al., 2016; Chung et al., 2017). It was shown that, in the zebrafish pronephros, proximo-distal segmentation was regulated by retinoic acid signaling rather than by Notch signaling (Liu et al., 2007; Wingert et al., 2007). It has yet to be determined if retinoic acid signaling regulates the mammalian nephron in a similar manner.

It is now well established that Wnt/β-catenin signaling is required for the maintenance and early differentiation of MNPs (Karner et al., 2011; Kuure et al., 2007; Park et al., 2012; Park et al., 2007; Ramalingam et al., 2018). Several studies have shown that Wnt/β-catenin signaling is also important at later stages of nephron development. When *Ctnnb1*, the gene encoding β-catenin, was deleted using *Pax8Cre* which targets both the developing nephron and CD in the mouse kidney, formation of the Bowman’s capsule was defective (Grouls et al., 2012). In addition, based on the pharmacological manipulation of Wnt/β-catenin signaling in mouse kidney explants, it was proposed that Wnt/β-catenin signaling promotes the formation of distal segments of the nephron and represses the formation of proximal segments (Lindstrom et al., 2014). Taken together, these studies suggest that Wnt/β-catenin signaling may play multiple roles in mammalian nephron segmentation.

In order to investigate how Wnt/β-catenin signaling regulates mammalian nephron patterning, we have performed genetic analyses of Wnt/β-catenin signaling by specifically targeting the developing nephron in the mouse kidney. Here, we report that epithelial nephron progenitor cells lacking β-catenin can form presumptive PT cells but cannot form differentiated PT cells. We also find that β-catenin is required for the formation of DT. Taken together, our data suggest that Wnt/β-catenin signaling regulates the development and maturation of multiple nephron segments in the mammalian kidney.

## Results

### Lineage analysis with *Osr2Cre* in the developing mouse kidney

In order to investigate the role of Wnt/β-catenin signaling in mammalian nephron patterning, we set out to perform β-catenin loss-of-function (LOF) study, specifically targeting the developing nephron in the mouse kidney. Since β-catenin is ubiquitously expressed in the kidney (Adam et al., 2017), the specificity of Cre is important. *Wnt4GFPcre* (*Wnt4*^*tm3(EGFP/cre)Amc*^ or *Wnt4*^*tm2(EGFP/cre)Svo*^) (Mugford et al., 2009; Shan et al., 2009) and *Pax8Cre* (*Pax8*^*tm1(cre)Mbu*^) (Bouchard et al., 2004) are widely used to target nephron tubules. However, the expression of these Cre lines is not exclusive to the developing nephron tubules; *Wnt4GFPcre* also targets the medullary stroma (Dirocco et al., 2013) and a subset of MNPs (Brunskill et al., 2014; Chung et al., 2017) while *Pax8Cre* targets the collecting duct in addition to the nephron lineage (Pietila et al., 2011). Removal of β-catenin from these non-nephron tubule cells may indirectly affect nephron patterning. Therefore, we chose to use *Osr2Cre* (*Osr2*^*tm2(cre)Jian*^) which was previously shown to specifically target developing nephrons in the mouse kidney (Lan et al., 2007). To examine which nephron segments are targeted by *Osr2Cre*, we performed lineage analysis employing Cre-mediated activation of a Rosa reporter (Ai3, *Gt(ROSA)26Sor*^*tm3(CAG-EYFP)Hze*^) (Madisen et al., 2010). We found that, in the SSB, *Osr2Cre* targeted the proximal and medial segments but not the distal segment (Figure 1A). To determine which nephron segments these Rosa reporter-positive cells in the SSB develop into, we performed co-immunostaining of EYFP and nephron segmentation markers. We found that the Rosa reporter was active in the podocytes, the Bowman’s capsule, PT, and LOH (Figure 1, A-D) but that it was inactive in the DT (Figure 1B). Furthermore, we found that, unlike *Wnt4GFPcre* or *Pax8Cre, Osr2Cre* did not target MNPs, medullary stroma, or the CD (Figure 1E). These data showed that *Osr2Cre* specifically targets all nephron segments except for the DT.

**Figure 1.**
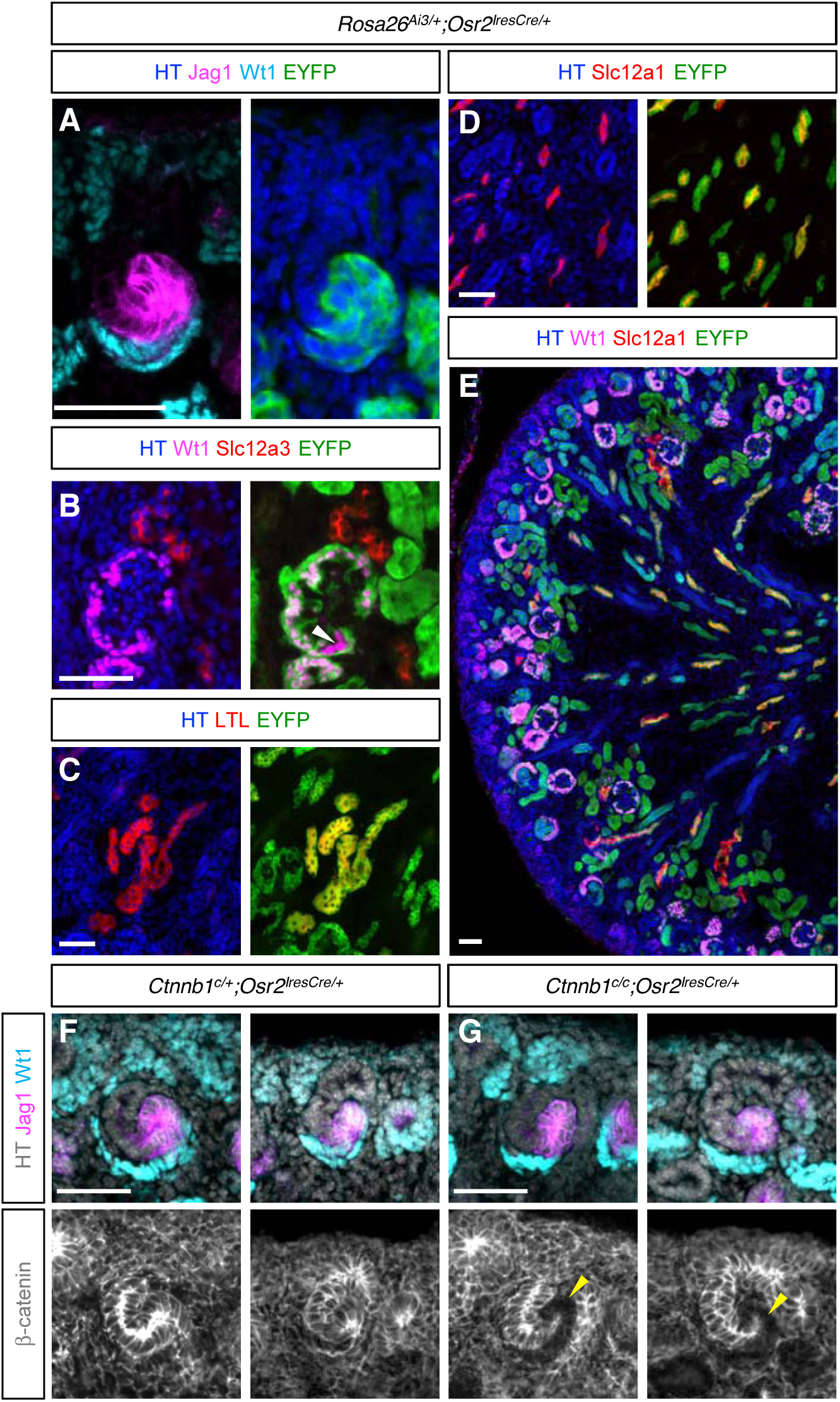
(A-E) Lineage analysis with *Osr2Cre* in the developing mouse kidney. (A) *Osr2Cre* targets the proximal (Wt1+) and medial (Jag1+) segments of the S-shaped body. In the nephron, *Osr2Cre* targets podocytes (B), proximal tubules (C), and loops of Henle (D), but not distal tubules (B). White arrowhead in (B) points to podocytes that escaped *Osr2Cre*. (E) *Osr2Cre* targets neither the cap mesenchyme nor the interstitial cells. (F) β-catenin is ubiquitously expressed in the S-shaped body. (G) *Osr2Cre* successfully removes β-catenin from the proximal and medial segments of the S-shaped body. The cells lacking β-catenin are marked with yellow arrowheads. HT, Hoechst. Stage E18.5. Scale bar: 50mm.

Next, we generated the β-catenin LOF mutant kidney with *Osr2Cre* and examined the presence of β-catenin in the SSB. In the control kidney, β-catenin was ubiquitously expressed in the SSB (Figure 1F). Consistent with *Osr2Cre*-mediated activation of the Rosa reporter in the SSB (Figure 1A), we found that *Osr2Cre* removed β-catenin from the proximal and medial segments of the SSB in the β-catenin LOF mutant kidney (Figure 1G). This allowed us to investigate the role of β-catenin in the development of the proximal nephron segments including, podocytes, PT, and LOH.

### β-catenin is required for the proper formation of the renal corpuscle

It was previously reported that the developing nephron lacking β-catenin failed to properly form the Bowman’s capsule (Grouls et al., 2012). However, it is unknown how loss of β-catenin affects the development of other nephron segments. To address this, we examined nephron segmentation markers in the β-catenin LOF mutant kidney with *Osr2Cre* (*Ctnnb1*^*c/c*^*;Osr2*^*IresCre/+*^) and its control (*Ctnnb1*^*c/+*^*;Osr2*^*IresCre/+*^) kidney. We used Wt1, *Lotus tetragonolobus* lectin (LTL), and Slc12a1 to mark podocytes, PT, and LOH, respectively. We found that all of these nephron segments were present in the mutant kidney (Figure 2, A-C), suggesting that the initial specification of these nephron segments were normal in the absence of β-catenin. To confirm that β-catenin was removed from each nephron segment in the mutant kidney, we examined the expression of β-catenin along with nephron segmentation markers. We found that, in the mutant kidney, these nephron segmentation marker-positive cells showed little to no β-catenin signal compared to their counterparts in the control kidney (Figure 2, A-C), showing that the β-catenin gene (*Ctnnb1*) was indeed deleted by *Osr2Cre*. These results suggest that β-catenin is dispensable for the initial specification of podocytes, PT, and LOH during nephron patterning. We also found that Slc12a3+ DT cells also were present in the mutant kidney (Figure 2D) but, since the DT cells were not targeted by *Osr2Cre* (Figure 1B), these cells were still positive for β-catenin (Figure 2D).

**Figure 2.**
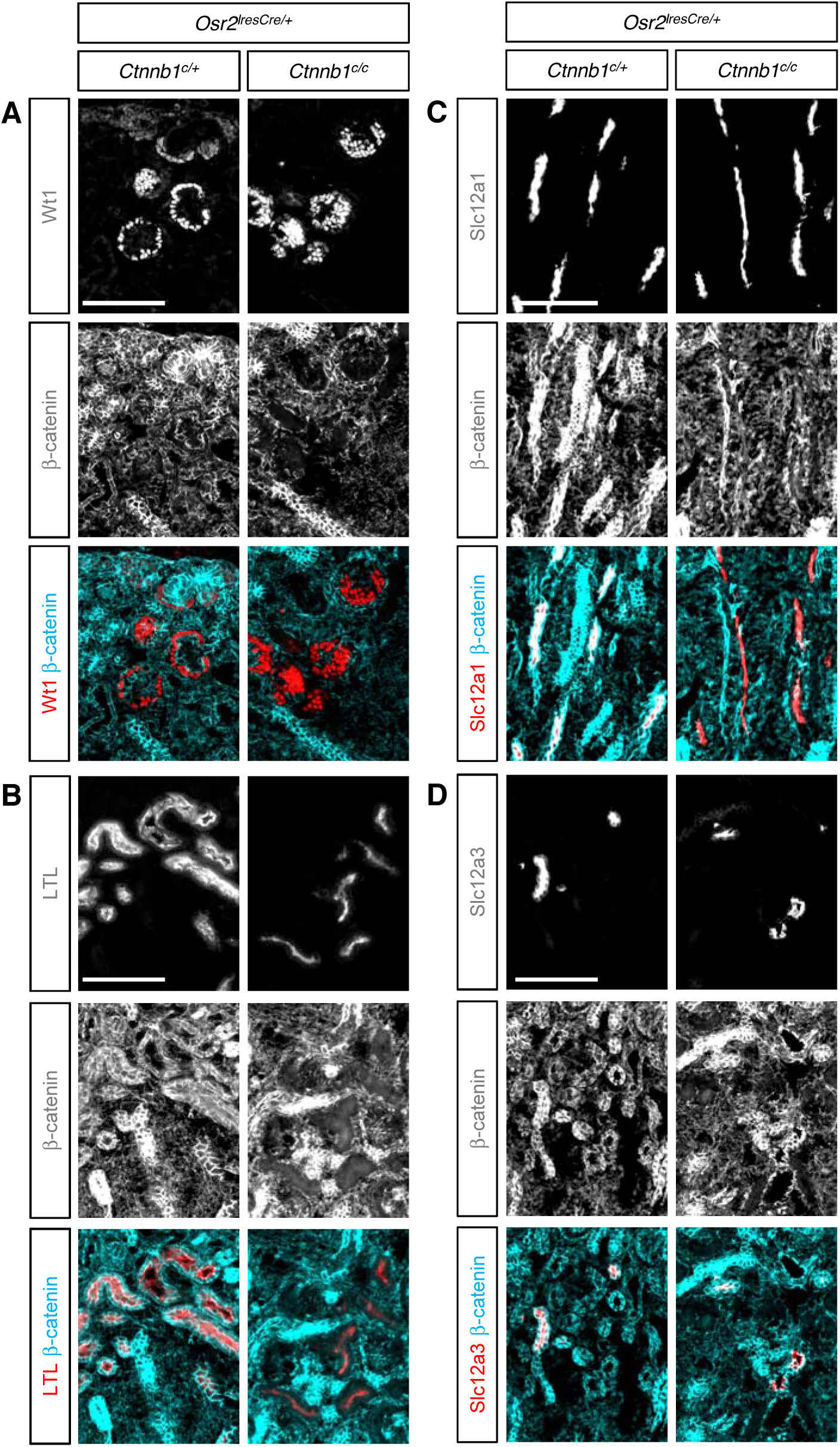
(A-C) In the β-catenin loss-of-function mutant kidney by *Osr2Cre*, the podocytes, proximal tubules, and loops of Henle show little or no β-catenin staining, suggesting that *Ctnnb1*, the gene encoding β-catenin, was deleted by *Osr2Cre.* Cells lacking β-catenin are able to form Wt1+ podocytes (A), LTL+ proximal tubules (B), and Slc12a1+ loops of Henle (C). (D) In the β-catenin mutant kidney, β-catenin is still present in Slc12a3+ distal tubules, consistent with the fact that *Osr2Cre* does not target distal tubules. Stage E18.5. Scale bar: 100mm.

Grouls *et al.* showed that, when β-catenin was removed by *Pax8Cre*, a subset of parietal epithelial cells in the Bowman’s capsule was replaced by podocytes, proposing that parietal epithelial precursor cells switch to the podocyte cell fate (Grouls et al., 2012). Consistent with their report, we observed a defective formation of the renal corpuscle in the β-catenin LOF mutant kidney by *Osr2Cre*. In the control kidney, Wt1+ podocytes surrounded Pecam1+ endothelial cells in the renal corpuscle (Figure 3A). However, in the β-catenin mutant kidney, endothelial cells failed to populate inside the renal corpuscle and podocytes failed to form the single cell layered crescent configuration characteristic of a normal renal corpuscle (Figure 3A). To assess if the aberrant formation of the renal corpuscle seen in the β-catenin LOF mutant kidney by *Osr2Cre* is caused by a defect in the early specification of visceral epithelial cells (presumptive podocyte progenitors) or parietal epithelial cells (presumptive Bowman’s capsule progenitors), we examined the SSB stage. We found that, in both control and mutant SSBs, endothelial cells migrated into the vascular cleft which was formed by the proximal and medial segments of the SSB, suggesting that β-catenin is dispensable for the visceral epithelial cells to recruit endothelial cells (Figure 3B). In the proximal segment of the SSB, the visceral and parietal epithelial cells showed distinct cell morphologies and differential gene expression. The visceral epithelial cells were organized in a columnar manner while the parietal epithelial cells were arranged in a thin end-to-end pattern (Figure 3B). Furthermore, the visceral epithelial cells expressed *Wt1* and *Mafb*, two genes required for podocyte development (Dong et al., 2015; Moriguchi et al., 2006; Sadl et al., 2002), while the parietal epithelial cells expressed *Wt1*, but not *Mafb* (Figure 3B). We found that the visceral and parietal epithelial cells lacking β-catenin showed normal cell morphologies and gene expression in the SSB (Figure 3B). Our results show that β-catenin is dispensable for the initial specification of visceral and parietal epithelial cells in the SSB.

**Figure 3.**
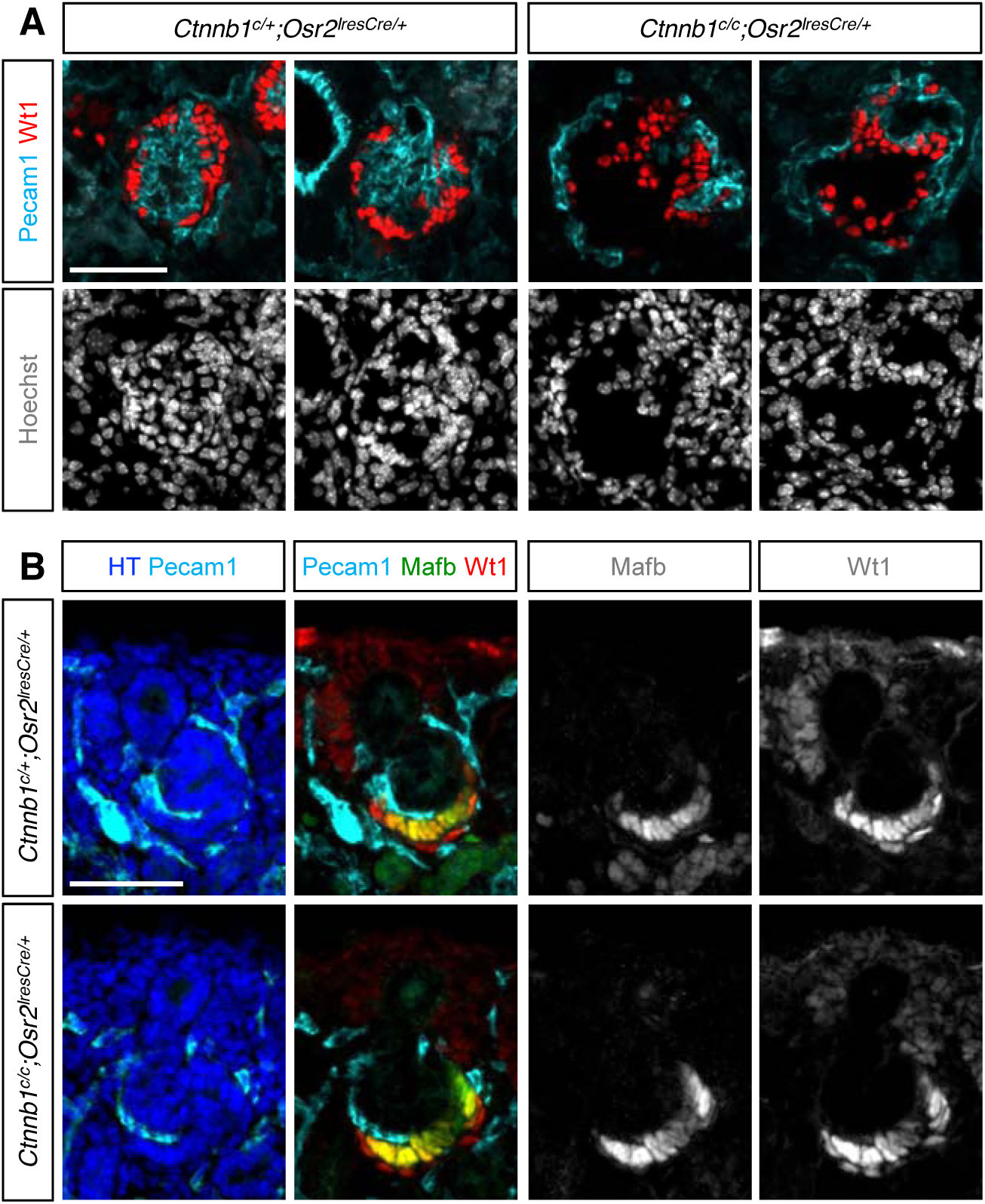
β-catenin is required for the proper formation of the renal corpuscle. (A) In the renal corpuscle of the control kidney, Wt1+ podocytes surround Pecam1+ endothelial cells in a crescent configuration. By contrast, in the β-catenin mutant kidney, Pecam1+ endothelial cells fail to populate inside the renal corpuscle. Two representative images from control and mutant are shown. (B) Pecam1+ endothelial cells invade the vascular cleft in the S-shaped body in both control and mutant kidneys. In the proximal segment of the S-shaped body, Wt1 is detected in both visceral and parietal epithelial cells while Mafb is detected in the visceral epithelial cells only. The initial specification of visceral and parietal epithelial cells appears normal in the β-catenin mutant kidney. HT, Hoechst. Stage E18.5. Scale bar: 50mm.

### β-catenin is required for presumptive proximal tubules to develop into differentiated proximal tubules

We found that the PT cells present in the β-catenin LOF mutant kidney showed substantially weaker LTL staining compared to the PT cells found in the control kidney (Figure 2B). Previously, we have shown that, in the developing mouse kidney, presumptive PT cells show weak LTL staining while differentiated PT cells show strong LTL staining and that presumptive PT cells lacking *Hnf4a* fail to develop into differentiated PT cells (Marable et al., 2018). Since weak LTL staining of PT cells in the β-catenin LOF mutant kidney was reminiscent of the phenotype seen in the *Hnf4a* mutant kidney, we examined if presumptive PT cells in the β-catenin LOF mutant kidney failed to develop into differentiated PT cells. It has been reported that *Cdh6* is expressed in presumptive PT cells and that its expression is downregulated in LTL-stained PT cells (Cho et al., 1998). Consistent with this, we found an inverse correlation between *Cdh6* expression and LTL staining in the control kidney (Figure 4). In addition, we found that both Cdh6-positive cells and LTL-stained cells expressed Hnf4a, a transcription factor specifically expressed in the PT cells (Lee et al., 2015) and required for PT development (Marable et al., 2018). Downregulation of *Cdh6* was accompanied by robust LTL staining and increased distance between Hnf4a+ nuclei, suggesting the maturation and enlargement of PT cells (Figure 4). Collectively, these observations suggest that Cdh6 is an excellent marker for presumptive PT cells. We found that, in the β-catenin LOF mutant kidney, most of the Hnf4a+ cells showed strong Cdh6 signal, indicating that they are presumptive PT cells (Figure 4). Cells with strong LTL staining were rarely found in the mutant kidney, suggesting that presumptive PT cells lacking β-catenin fail to develop into differentiated PT cells (Figure 4). Our results show that Wnt/β-catenin signaling is required for presumptive PT cells to further develop into differentiated PT cells.

**Figure 4.**
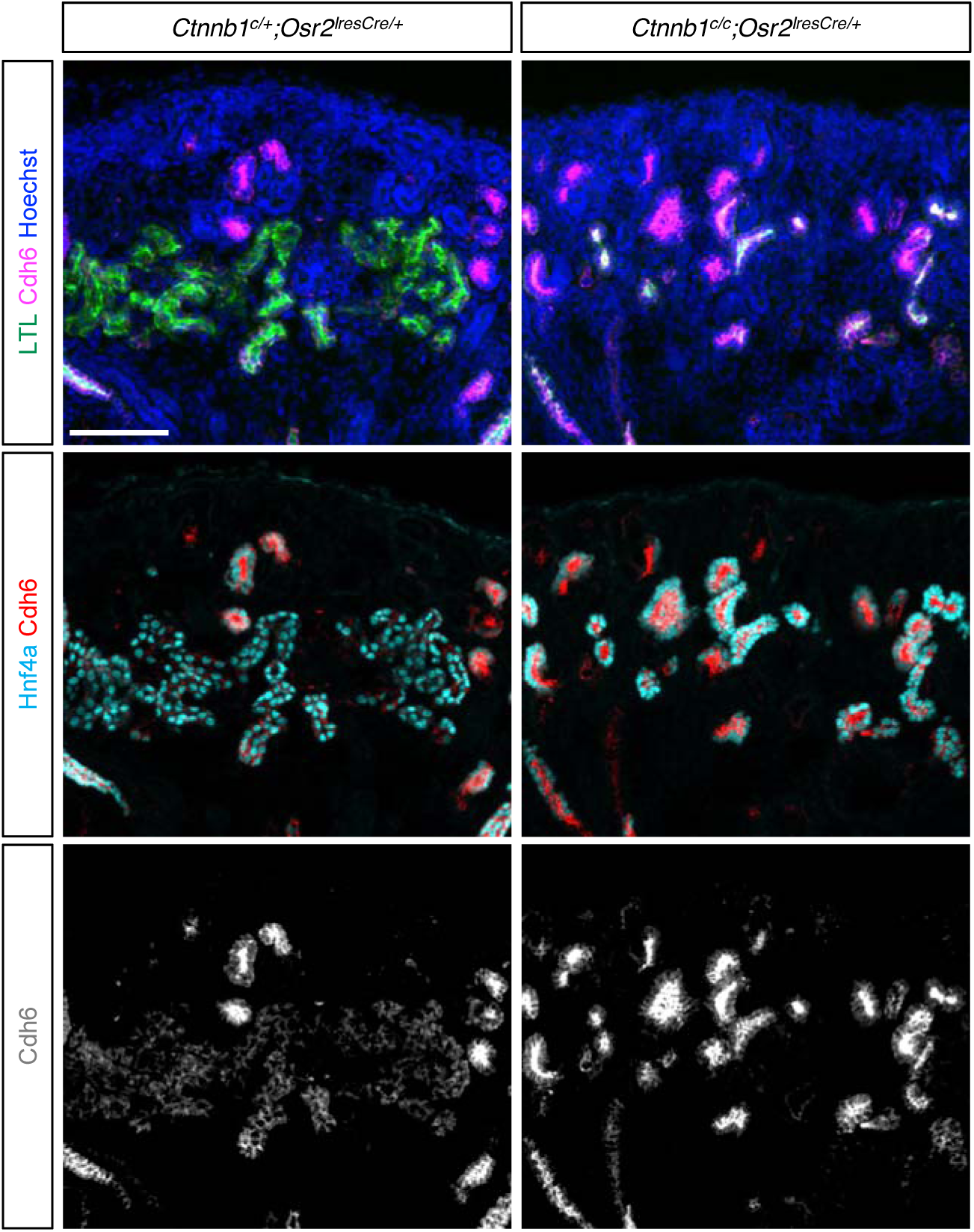
β-catenin is required for the formation of differentiated proximal tubule cells with strong LTL staining. Hnf4a marks both presumptive and differentiated proximal tubules. In the control kidney, presumptive proximal tubules show strong Cdh6 signal and weak LTL staining while differentiated proximal tubules show weak Cdh6 signal and strong LTL staining. In the β-catenin loss-of-function mutant kidney by *Osr2Cre*, all Hnf4a+ cells show strong Cdh6 signal, failing to form differentiated proximal tubules with strong LTL staining. Stage E18.5. Scale bar: 100mm.

### β-catenin is required for the formation of the distal tubules

Since *Osr2Cre* does not target the DT cells, it does not allow us to investigate how Wnt/β-catenin signaling regulates the formation of DT. To address this, we generated the β-catenin LOF mutant kidney (*Ctnnb1*^*c/c*^*;Wnt4*^*GFPcre/+*^) using *Wnt4GFPcre* (*Wnt4*^*tm3(EGFP/cre)Amc*^) (Mugford et al., 2009). We have previously shown that *Wnt4GFPcre* can target the nephron lineage including DT (Chung et al., 2017). Considering the fact that *Wnt4GFPcre* also targets the medullary stroma (Dirocco et al., 2013), *Wnt4GFPcre* may not be suitable for studying the role of β-catenin in LOH since removal of β-catenin in the medullary stroma may affect the development of the adjacent LOH. However, this is less of a concern for studying the DT because the DT cells are located close to the cortical region of the developing mouse kidney, apart from the medullary stroma.

Consistent with our observation that PT development was arrested in the β-catenin LOF mutant kidney by *Osr2Cre* (Figure 4), the β-catenin LOF mutant kidney by *Wnt4GFPcre* also showed fewer PT cells and weaker LTL staining (Figure 5A). In the control kidney, Slc12a1+ cells elongated toward the papilla region, forming the characteristic LOH structure. By contrast, in the β-catenin LOF mutant kidney by *Wnt4GFPcre*, Slc12a1+ cells failed to elongate, stunting the development of the papilla (Figure 5A). Considering the fact that the β-catenin LOF mutant kidney by *Osr2Cre* showed relatively normal LOH formation (Figure 2A), the failed elongation of LOH seen in Figure 5A was likely to be caused by removal of β-catenin in the medullary stroma by *Wnt4GFPcre*.

**Figure 5.**
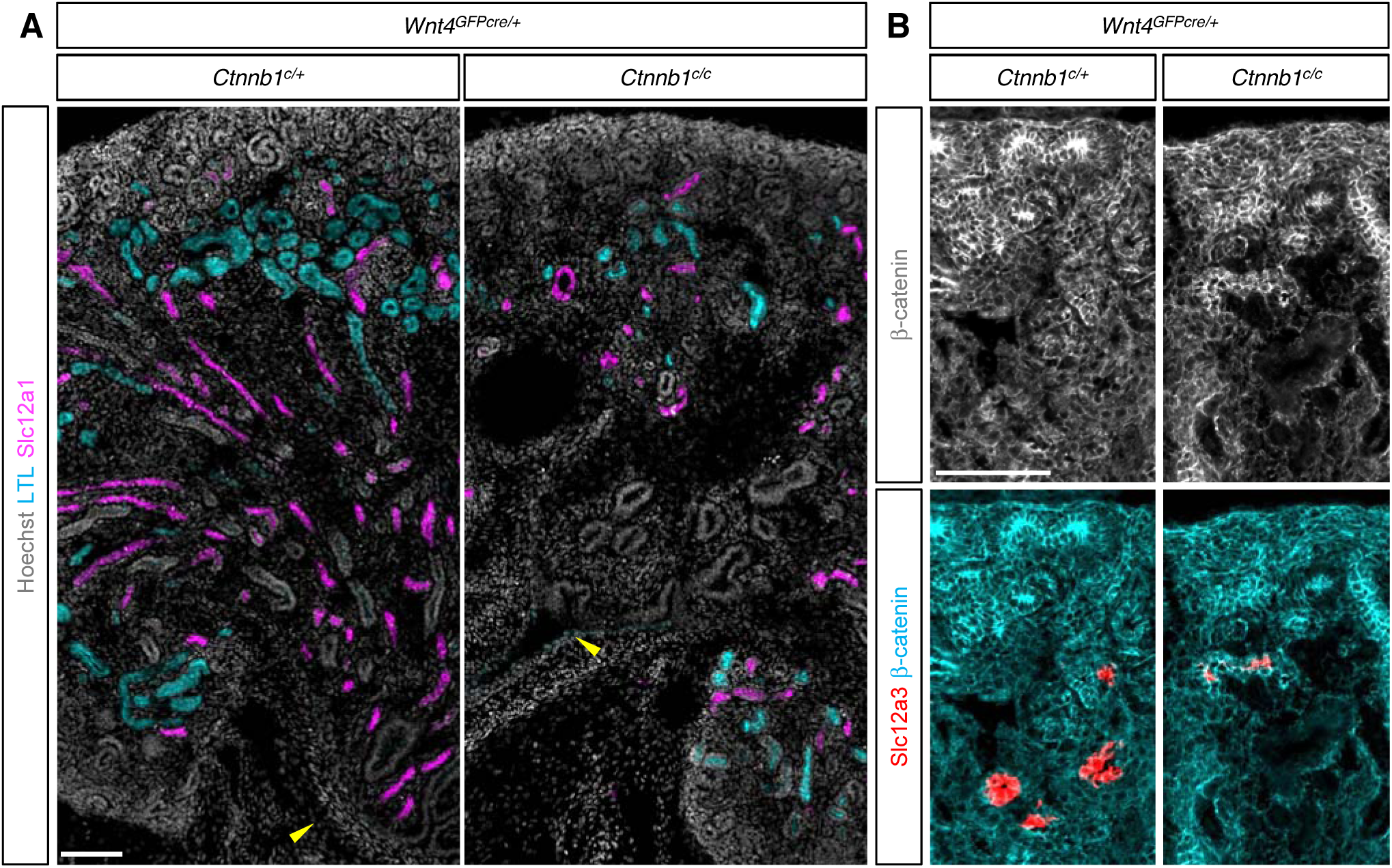
Removal of β-catenin by *Wnt4Cre* inhibits loop of Henle elongation and distal tubule formation. (A) Slc12a1+ cells in the control kidney elongate, contributing to papilla formation. Slc12a1+ cells in the β-catenin loss-of-function mutant kidney by *Wnt4Cre* fail to elongate, causing a defective papilla formation. The ureter is marked by yellow arrowheads. (B) In the β-catenin loss-of-function mutant kidney by *Wnt4Cre*, all Slc12a3+ distal tubules are positive for β-catenin, suggesting that these cells have escaped *Wnt4Cre*-mediated removal of β-catenin. The absence of β-catenin-negative distal tubule cells suggests that β-catenin is required for the formation of distal tubules. Stage E18.5. Scale bar: 100mm.

When *Ctnnb1* was deleted with *Wnt4GFPcre*, we found that the mutant kidney formed the DT as marked by Slc12a3 (Figure 5B), a DT-specific transporter (Lee et al., 2015). However, all DT cells found in the mutant kidney were positive for β-catenin (Figure 5B), suggesting that these cells had escaped *Wnt4GFPcre*-mediated removal of β-catenin. The absence of β-catenin-negative DT cells in the mutant kidney suggests that Wnt/β-catenin signaling is required for the formation of the DT.

### Constitutive activation of Wnt/β-catenin signaling in the developing nephron blocks proper nephron segmentation

Our β-catenin LOF studies suggest that Wnt/β-catenin signaling regulates the development of multiple nephron segments. To further explore the mechanism of Wnt/β-catenin signaling in nephron patterning, we performed a β-catenin gain-of-function (GOF) study using *Osr2Cre*. The conditional allele used in this study has exon3 of the *Ctnnb1* gene flanked by two *LoxP* sites (*Ctnnb1*^*ex3*^) (Harada et al., 1999). Since exon3 encodes the N-terminus of β-catenin protein, Cre-mediated recombination results in the production of an N-terminally truncated β-catenin which is resistant to degradation. Increased abundance of β-catenin leads to the activation of canonical Wnt signaling in a ligand-independent manner (Harada et al., 1999).

We found that the β-catenin GOF mutant kidney by *Osr2Cre* lacked glomeruli and PTs (Figure 6). Slc12a1+ LOH cells and Slc12a3+ DT cells were present in the β-catenin GOF mutant kidney but they failed to elongate or form normal LOH or DT segments (Figure 6). We found no evidence that constitutive activation of Wnt/β-catenin signaling in the developing nephrons promoted the formation of any specific nephron segment. Instead, we found that expression of a stabilized form of β-catenin in the developing nephron inhibited the formation of all nephron segments.

**Figure 6.**
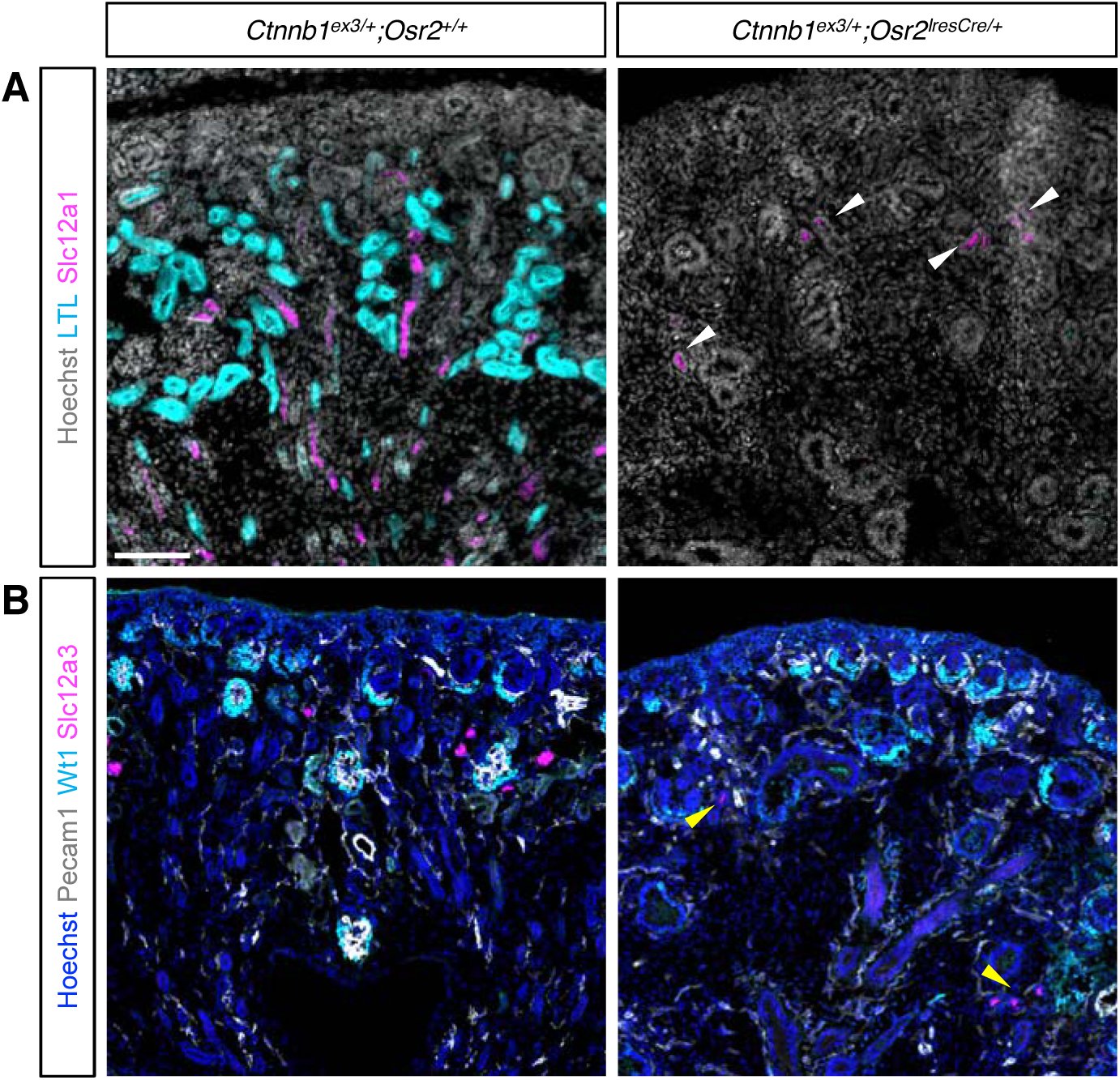
Expression of a stabilized form of β-catenin inhibits proper nephron patterning. The β-catenin gain-of-function mutant kidney by *Osr2Cre* fail to form properly patterned nephrons. No proximal tubules (A) or glomeruli (B) are formed. Slc12a1+ loop of Henle (marked by white arrowheads, A) or Slc12a3+ distal tubule (marked by yellow arrowhead, B) cells are present but they fail to elongate. Stage E18.5. Scale bar: 100mm.

To investigate how constitutive activation of Wnt/β-catenin signaling interferes with nephron patterning, we examined early development of the nephron in the β-catenin GOF mutant kidney. In the control kidney, the proximal and medial segments of the SSB were marked by Wt1 and Jag1, respectively, and the expression domains of Wt1 and Jag1 did not overlap, showing clear separation (Figure 7A). By contrast, in the β-catenin GOF mutant kidney, we observed an overlap of the Wt1 expression domain and the Jag1 expression domain (white arrowhead, Figure 7A). Although the initial morphology of the SSB appeared normal in the β-catenin GOF mutant kidney, the Jag1 expression domain expanded aberrantly toward the Wt1 expressing proximal segment of the SSB (yellow arrowhead, Figure 7A). This result shows that the β-catenin GOF mutant kidney by *Osr2Cre* exhibits a defect in nephron segmentation as early as the SSB stage.

**Figure 7.**
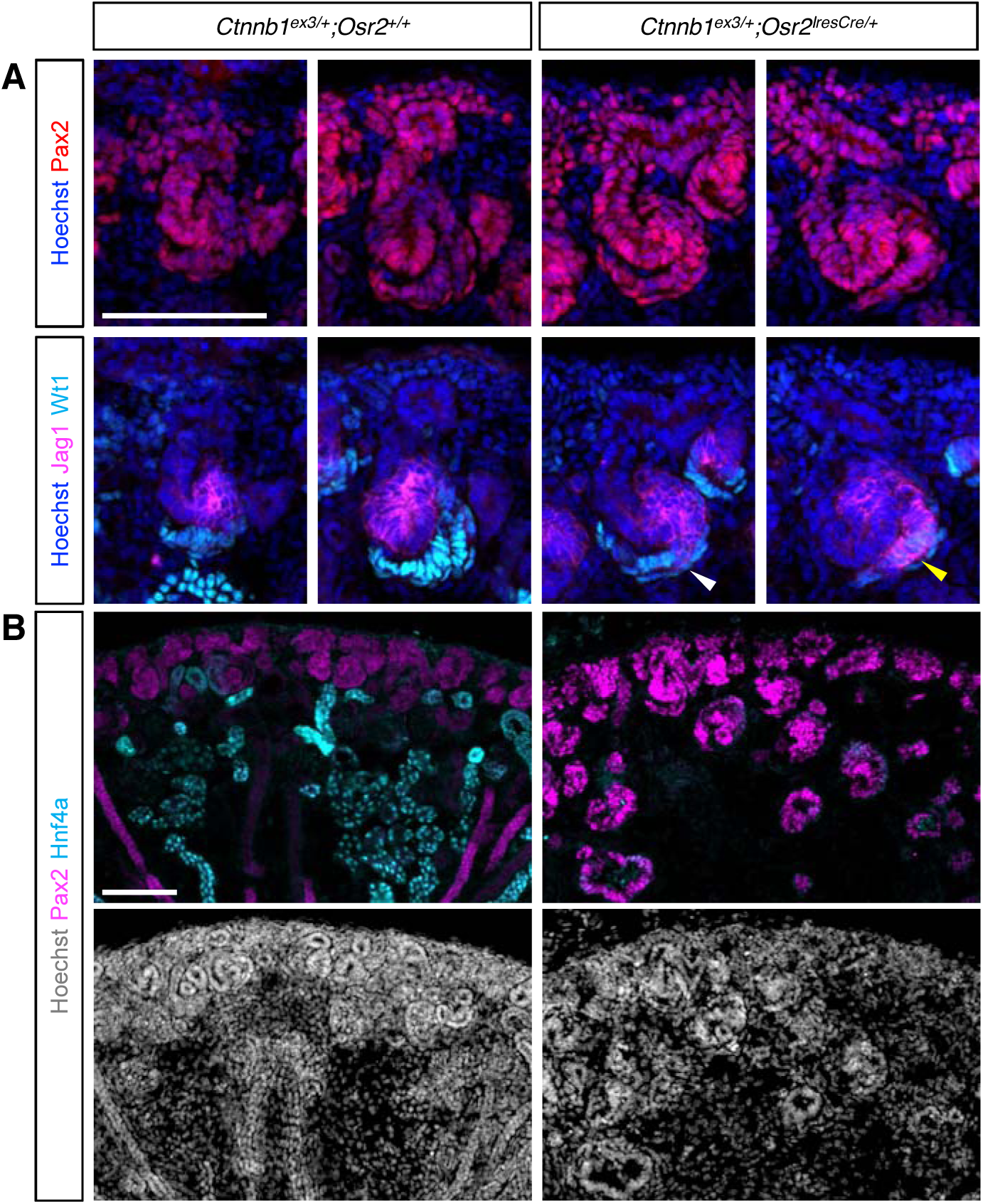
Expression of a stabilized form of β-catenin keeps epithelial nephron progenitors from further differentiating. (A) In the control kidney, Jag1 and Wt1 mark the medial and proximal segments of the S-shaped body, respectively. In the β-catenin gain-of-function mutant kidney by *Osr2Cre*, the S-shaped body is formed initially (marked by white arrowhead) but the Jag1 expression domain invades into the Wt1 expression domain (marked by yellow arrowhead). Pax2 marks the collecting duct, the cap mesenchyme, and nascent developing nephrons. (B) In the control kidney, Pax2 expression in epithelial nephron progenitors is downregulated after S-shaped body stage. As a result, most of the Hnf4a+ cells are negative for Pax2. In the β-catenin gain-of-function mutant kidney by *Osr2Cre,* Pax2 is ectopically expressed in the nephron lineage and little or no Hnf4a expression is detected, suggesting that β-catenin gain-of-function mutant cells fail to exit from their progenitor status. Stage E18.5. Scale bar: 100mm.

One of the most striking defects seen in the β-catenin GOF mutant kidney by *Osr2Cre* was the complete lack of LTL+ PTs (Figure 6A). Considering the fact that Hnf4a plays a critical role in PT development (Marable et al., 2018), we examined Hnf4a expression in the β-catenin GOF mutant kidney. In the nephron lineage of the control kidney, Pax2 expression was mostly restricted to the nephrogenic zone at the cortex, marking the cap mesenchyme and nascent developing nephrons (Figure 7B). Pax2 was undetectable in most of the Hnf4a+ PT cells, suggesting that the formation of PTs is accompanied by downregulation of Pax2 (Figure 7B). Interestingly, we found no Hnf4a expression in the β-catenin GOF mutant kidney (Figure 7B), suggesting that the lack of PTs in the mutant kidney is caused by failure to activate Hnf4a expression. Pax2 expression was not restricted to the nephrogenic zone in the mutant kidney. Instead, most of the epithelial structures found in the mutant kidney showed persistent expression of Pax2 (Figure 7B), suggesting that epithelial nephron progenitors experiencing constitutive activation of Wnt/β-catenin signaling failed to further differentiate into PTs. Taken together, our results suggest that tight regulation of Wnt/β-catenin signaling is required for the proper patterning of the nephron even after initial commitment of nephron progenitors.

## Discussion

Wnt/β-catenin signaling is known to regulate the maintenance and commitment of the MNPs (Karner et al., 2011; Kuure et al., 2007; Park et al., 2012; Park et al., 2007; Ramalingam et al., 2018). Here we show that it continues to play critical roles in nephron patterning after the initial commitment of nephron progenitors. Strikingly, we found that Wnt/β-catenin signaling regulates the development of multiple nephron segments without promoting the formation of any specific nephron segment.

Little is known about how multipotent MNPs develop into different nephron segments during kidney development (Desgrange and Cereghini, 2015). The first sign of nephron segmentation can be seen as early as the RV stage (Cho et al., 1998; Dressler, 2006; Georgas et al., 2009; Mugford et al., 2009). The distal part of the RV (the part closer to the branching tip of the collecting duct) shows differential gene expression compared to the rest of the RV. We have previously identified multiple β-catenin-bound enhancers that are associated with genes activated during differentiation of MNPs (Park et al., 2012). Transgenic analyses in the developing mouse kidney revealed that these enhancers were active in the distal RV but inactive in the proximal RV (Park et al., 2012). In addition, we have previously shown that the expression of *Jag1* in the distal RV was correlated with the downregulation of *Six2* while Six2 was still detectable in the proximal RV (Chung et al., 2016). The cellular identity of the MNPs appears closer to that of the proximal RV than that of the distal RV, consistent with a recent finding that the cells in the proximal RV exit from the cap mesenchyme later than the cells of the distal RV (Lindstrom et al., 2018).

By the time the RV is transformed into the SSB, three discrete (proximal, medial and distal) segments are established. The proximal segment of the SSB is thought to develop into the renal corpuscle because it expresses podocyte marker genes such as *Wt1* and *Mafb* (Figure 3B). The distinct cell morphologies of the two cell layers in this segment suggest that the visceral and parietal epithelial cells develop into podocytes and Bowman’s capsule, respectively. The fact that *Osr2Cre* targets the medial segment, but not the distal segment, of the SSB helped us reveal that the medial segment develops into PT and LOH. This is consistent with our previous report that Hnf4a, a transcription factor required for the PT development, was detected in the proximal subset of the medial segment of the SSB (Marable et al., 2018) and suggests that Hnf4a-negative cells in the distal subset of the medial segment develop into LOH. This then allowed us to infer that the distal segment of the SSB develops into the DT. Our lineage analysis with *Osr2Cre* uncovered a correlation between SSB segments and mature nephron segments.

Since the β-catenin LOF mutants by *Osr2Cre* die shortly after birth with severe craniofacial defects (Chen et al., 2009), we could not analyze the mutant kidney postnatally. By contrast, the β-catenin LOF mutants by *Pax8Cre* show no lethality until 2 weeks of age (Grouls et al., 2012). In the kidney, *Pax8Cre* targets DT and CD in addition to the nephron segments targeted by *Osr2Cre*. Previous electron microscopy analysis of the β-catenin LOF mutant kidney by *Pax8Cre* showed a glomerulogenesis defect similar to our results shown in Figure 3A. Furthermore, it was reported that the β-catenin LOF mutant adult kidney by *Pax8Cre* lacks the outermost cortex (Grouls et al., 2012), implying a similar PT defect seen in our study. However, the effects on other nephron segments were not investigated in that report.

We have previously shown that, in the absence of *Hnf4a*, presumptive PT cells failed to develop into differentiated PT cells, suggesting that Hnf4a is dispensable for the formation of presumptive PT cells but required for the formation of differentiated PT cells (Marable et al., 2018). It is quite interesting that removal of β-catenin prevented presumptive PT cells from developing into differentiated PT cells, showing a similar defect seen in the *Hnf4a* LOF mutant kidney. A number of reports suggested a potential interaction between the Wnt/β-catenin pathway and Hnf4a (Gougelet et al., 2014; Neve et al., 2014; Norton et al., 2014; Vuong et al., 2015; Yang et al., 2013). Notably, it was shown that Hnf4a and β-catenin not only form a complex but also share common target genes in the liver (Gougelet et al., 2014). Identification of common target genes regulated by both Hnf4a and β-catenin in presumptive PT cells would help us understand how these two factors coordinate PT development.

We found that the β-catenin LOF mutant kidney developed structures resembling LOH as judged by the expression of *Slc12a1*, a gene encoding a LOH-specific transporter. However, the mutant LOHs were substantially thinner compared to those found in the control kidney (Figure 2C). Therefore, it is possible that Wnt/β-catenin signaling regulates the maturation of LOH. Since *Osr2Cre* does not target the DT cells, we generated another β-catenin LOF mutant kidney using *Wnt4GFPcre* which does target DT (Chung et al., 2017). We found that all the DT cells formed in the β-catenin LOF mutant kidney were positive for β-catenin (Figure 5), indicating that these cells escaped Cre-mediated removal of the β-catenin gene. The absence of β-catenin-negative DT cells in the mutant kidney suggests that β-catenin is required for the formation of DT. Our result is consistent with a previous report that the distal segment of the SSB is exposed to a higher level of Wnt/β-catenin signaling than the rest of the SSB (Lindstrom et al., 2014). This is also supported by the fact that the β-catenin-bound enhancers identified in MNPs are active in the distal segment of the SSB (Park et al., 2012). Although the distal segments of the developing nephron may experience a higher level of Wnt/β-catenin signaling, we found that constitutive activation of Wnt/β-catenin signaling did not promote the formation of the distal segments. Instead, it blocked the proper patterning of all nephron segments (Figure 6). Considering that the β-catenin GOF mutant cells show persistent expression of Pax2 and that they fail to activate Hnf4a expression (Figure 7), it appears that constitutive activation of Wnt/β-catenin signaling in epithelialized nephron progenitors results in developmental arrest.

Overall, our data suggest that Wnt/β-catenin signaling regulates the patterning of multiple nephron segments along the proximo-distal axis. It is likely that precise spatiotemporal activation and inactivation of Wnt/β-catenin signaling is required for the proper patterning of the mammalian nephron.

## Materials and Methods

### Mouse strains

*Gt(ROSA)26Sor*^*tm3(CAG-EYFP)Hze*^ (*Rosa26*^*Ai3*^) (Madisen et al., 2010), *Wnt4*^*tm3(EGFP/cre)Amc*^ (*Wnt4GFPcre*) (Mugford et al., 2009), *Osr2*^*tm2(cre)Jian*^ (*Osr2Cre or Osr2*^*IresCre*^) (Lan et al., 2007), *Ctnnb1*^*tm2Kem*^ (*Ctnnb1*^*c*^) (Brault et al., 2001), and *Ctnnb1*^*tm1Mmt*^ (*Ctnnb1*^*ex3*^) (Harada et al., 1999) mice were described previously. All mice were maintained in the Cincinnati Children’s Hospital Medical Center (CCHMC) animal facility according to animal care regulations. The Animal Studies Committee at CCHMC approved the experimental protocols (IACUC2013-0105 and IACUC2017-0037). We adhere to the NIH Guide for the Care and Use of Laboratory Animals.

### Immunofluorescence

Embryonic kidneys at E18.5 were fixed with 4% paraformaldehyde in PBS at room temperature for 10 min, incubated in 10% sucrose in PBS at 4°C overnight, and imbedded in OCT. Cryosections of 10 or 12μm were incubated in PBS containing 0.1% Triton x-100, 5% heat-inactivated sheep serum, and the following antibodies: Jag1 (TS1.15H, rat, 1:20, DSHB), Wt1 (sc-7385, mouse IgG1, 1:100, Santa Cruz), GFP (GFP-1020, chick IgY, 1:500, Aves Labs), Slc12a3 (HPA028748, rabbit, 1:300, Sigma), LTL (FL-1321, FITC, 1:200, Vector Laboratories), Slc12a1 (18970-1-AP, rabbit, 1:200, Proteintech), β-catenin (71-2700, rabbit, 1:500, Invitrogen), β-catenin (sc-7963, mouse IgG1, 1:50, Santa Cruz), Pecam1 (sc-18916, rat, 1:300, Santa Cruz), Mafb (HPA005653, rabbit, 1:300, Sigma), Cdh6 (HPA007047, rabbit, 1:300, Sigma), Hnf4a (ab41898, mouse IgG2a, 1:500, Abcam), Pax2 (21385-1-AP, rabbit, 1:200, Proteintech). Fluorophore-conjugated secondary antibodies (Thermo Fisher Scientific or Jackson ImmunoResearch Laboratories) were used at a 1:500 dilution. Nuclei were stained with Hoechst 33342 before mounting. Images were acquired on a Nikon TiE microscope with Andor Zyla 4.2 camera and Lumencor SpectraX light source housed at the Confocal Imaging Core at CCHMC.

## Acknowledgments

We thank Steve Potter and Cristina Cebrian for critically reading our manuscript.

## Author contributions

Conceptualization: P.D., C.W.C., J.-S.P.; Methodology: P.D., E.C., J.-S.P.; Formal analysis: P.D., C.W.C., E.C., J.-S.P.; Investigation: P.D., C.W.C., E.C., J.-S.P.;

Resources: J.-S.P.; Data curation: P.D., C.W.C., E.C., J.-S.P.; Writing - original draft:

E.C., J.-S.P.; Writing - review & editing: E.C., J.-S.P.; Supervision: J.-S.P.; Project administration: J.-S.P.; Funding acquisition: J.-S.P.

